# A neurocomputational basis of face recognition changes in ASD: E/I balance, internal noise, and weak neural representations

**DOI:** 10.1101/2025.04.02.646903

**Authors:** Xijing Wang, Emily Rios, Lang Chen

**Affiliations:** Mathematics and Computer Science, Santa Clara University; Neuroscience Program, Santa Clara University; Psychology, Santa Clara University

**Keywords:** Key word: Face recognition, ASD, E/I imbalance, internal noise, CNN models

## Abstract

Individuals with Autism Spectrum Disorder (ASD) are known for their socio-communicative challenges, including face recognition. Despite mounting evidence in behavioral studies, the neurocomputational basis of these challenges remains unclear. Meanwhile, neurobiological theories propose that ASD may arise from an imbalance of excitatory and inhibitory signals (E/I imbalance) or excessive internal noise (IN). However, studies with humans can hardly provide causal evidence. Therefore, this study employed Conventional Neural Network (CNN) models to simulate face recognition in typical populations and ASD based on the claims of I/E imbalance and IN theories. By varying the positive slope in the ReLU activation function (simulating E/I imbalance) and random noises added to the weights (simulating internal noise), we showed that CNN models with non-optimal ReLU slope or noised weights led to poorer performance in face recognition and atypical neural representations of faces. Overall, simulations based on the E/I imbalance theory seem to encompass a broader range of behavioral profiles in ASD. Our approach to using CNN models to test neurobiological theories is highly theory-driven, and our results provided causal evidence to how neurobiological factors could influence face recognition in ASD. This framework could be easily adapted to test in other neurobiological disorders, providing a plausible bridge between neurobiological theories and behavioral and neuroimaging research on humans.

---

Autism Spectrum Disorder (ASD) is characterized by socio-communicative challenges^1^, but autistic individuals also show cognitive challenges in face recognition^2,3^, a crucial factor for social interaction^4^. Behavioral studies have suggested that autistic individuals demonstrate lower performance in face recognition and discrimination^5–8^, particularly when tasks involve memory^9^. Functional Magnetic Resonance Imaging (fMRI) studies have shown atypical neural activations or connectivity in autistic individuals during face processing^10–15^. Nevertheless, the underlying neurocomputational mechanism of atypical face recognition in ASD remains unclear^16^. In this study, therefore, we used a computational approach to the role of two neurobiological factors, namely, the imbalance of excitatory and inhibitory signals (E/I imbalance)^17^ and internal noise (IN)^18^ in disrupted face recognition in ASD.

The excitation/inhibition imbalance (E/I imbalance) theory hypothesizes that the imbalance ratio between excitatory (e.g., glutamatergic) and inhibitory (e.g., GABAergic) neural signals underlie many developmental disorders including ASD^17,19,20^. Indeed, neuroimaging studies have shown that, compared to the non-ASD group, autistic individuals show excessive activations to sensory stimuli^10,21^ and sometimes to face stimuli^12,14^ as well as aberrant functional connectivity ^22–25^ Another theory, known as internal noise (IN) theory, provides a different lens of the cognitive challenges observed in ASD, positing that autistic individuals experience higher levels of intrinsic noise, adversely affecting neural processing efficiency^18^. Specifically, the increased internal noise leads to unusually large fluctuations in neural responses, resulting in unreliable and less predictable representations of the environment, resulting in neurocognitive deficits in ASD^26,27.^ This theory is also supported by neuroimaging studies showing the variability of neural response in autistic individuals is higher than in the non-ASD group^27–29^. So far, it is challenging for neuroimaging studies on humans to reconcile the two different theories.

Face recognition is an ideal case to test the predictions of these two neurobiological theories. Human faces are highly similar and overlapping perceptually^30^, posing an intrinsic challenge to an imbalanced or noisy learning system. Indeed, previous neuroimaging studies have shown that autistic individuals showed hypoactivation in the core face processing area (such as the face fusiform area; FFA)^31^ but extensive activations in other brain regions^11,12,32^, suggesting a neurocomputational challenge to store and process the representations of face stimuli^33,34^. Furthermore, two neurobiological theories could have different predictions on the neural representations of face stimuli in ASD. According to the E/I imbalance theory, excessive excitatory signals could lead to diffused activations in the brain, resulting in undifferentiated neural representations and reduced ability for object recognition^35,36^. Overexcitation will likely result in increased similarity of neural representations for different persons (between-person similarity) but little neural representations of faces of the same person (i.e., within-person similarity). Meanwhile, the IN theory argues that increased internal noises in the brain will create unstable neural responses with random fluctuations, leading to weakened and unreliable neural representations of individual faces. As a result, both between-person and within-person similarity of neural representations should be reduced, hampering the ability to recognize faces. Thus, although not mutually exclusive, the two theories have different predictions of underlying neural representations that result in face recognition challenges in ASD.

In this study, we used computational models to test the predictions of the two theories, because they have been used to examine theories of cognitive and neurobiological disorders for a long time ^37–39^. Recent advances have demonstrated that computational models cannot only simulate cognitive behaviors and deficits but also provide a window to examine individual variabilities in internal representations^39,40^. Specifically, the Convolutional Neural Network (CNN) model is a dominant tool to approximate visual processing and object recognition^41,42^ with reasonable resemblance to the ventral visual pathway in humans^43^ with populational activations^44^, and processing hierarchy^44^. Studies have shown that CNN models can successfully explain the variance in neural responses in the inferotemporal cortex neurons for image recognition^45^, show high correlations with brain responses in the visual cortex of humans^46^, and demonstrate benchmark findings of face recognition behaviorally^47^. Therefore, we will use a theory-driven approach to simulate face recognition in CNNs, and manipulate key model parameters based on neurobiological theories (E/I imbalance and IN). By modulating in these biologically meaningful factors, we planned to demonstrate varying behavioral outcomes and internal representations for face recognition accordingly. In this way, we could test the neurobiological theories of ASD within well-controlled learning systems, i.e., CNN models, and provide causal evidence to elucidate potential neurocomputational mechanisms of face recognition challenges in ASD.

## Method

### General CNN model architecture

We implemented standardized Convolutional neural network (CNN) models^48,49^ with one input layer, three convolutional Blocks, a flattening Layer, and a dense Layer (i.e., the output layer). Each convolutional block consisted of convolutional, normalization, pooling (size = 2*2), and dropout (rate = 0.2) layers (**Figure 1A)**. In the last dense layer, the flattened output was processed by the SoftMax function to produce a probability value for the identity of each person (10 units, each for one person’s identity). The filter size of each convolutional layer was set to 16, 32, and 64 as practiced in the literature. When we tested the hypotheses of the E/I imbalance and IN theories in the CNN models, we manipulated two parameters: the slope of the ReLU function and the standard deviation of the Gaussian noises added to all the weights, respectively. We will explain the details in the training and testing procedure section.

**Figure 1.**
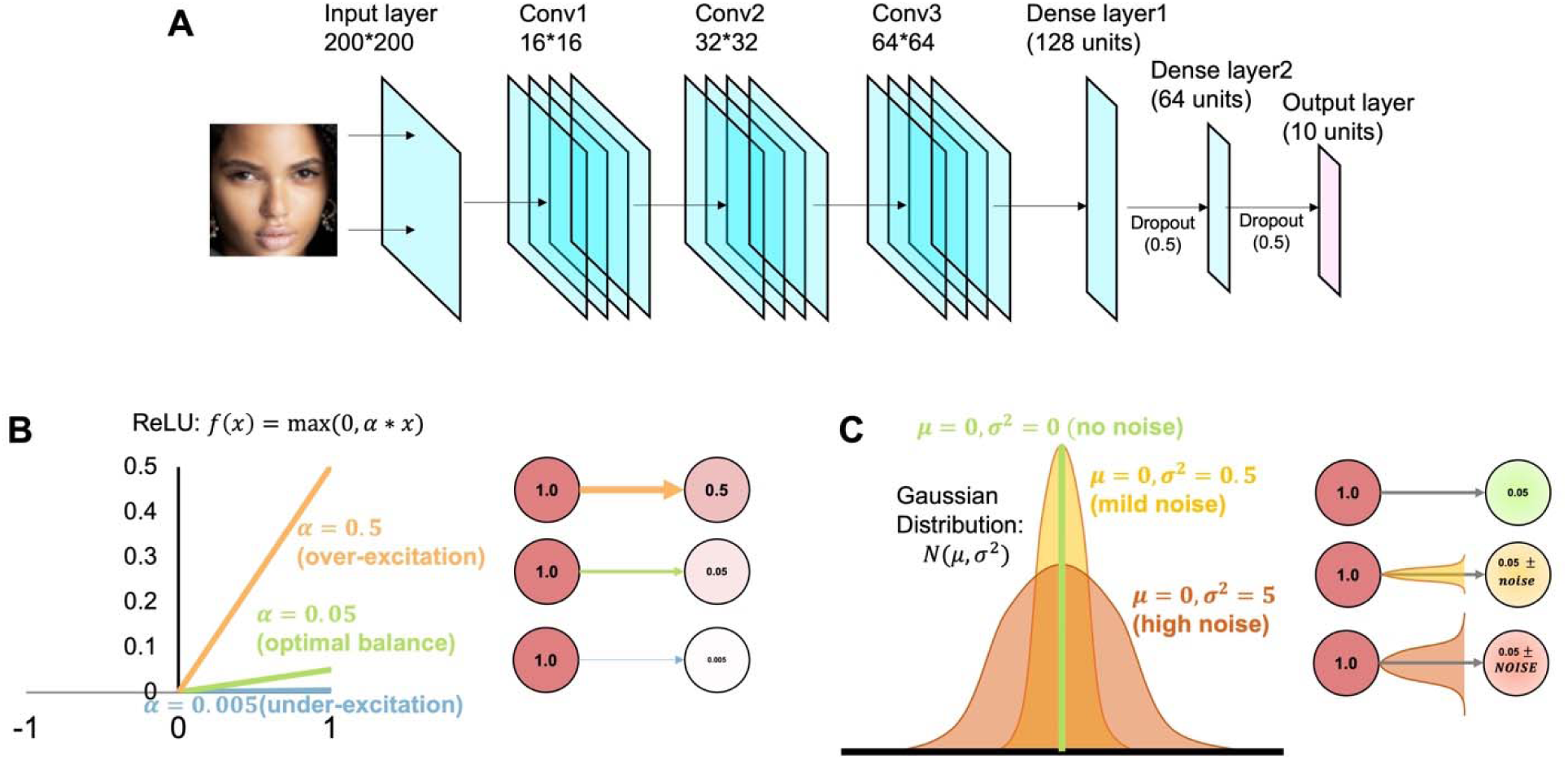
Illustrations of the architecture of the CNN models (**A**) and the manipulations of E/I imbalance (i.e., the slope of the ReLU function; **B**) and internal noise (i.e., Gaussian noise on weights; **C**).

### Training and testing sets

The training and testing set of face pictures in this study comprised 50 selected images from 10 celebrities, with five images representing each celebrity’s identity. All images were gathered from the internet, including 5 males and 5 females, as well as from diverse demographic backgrounds such as age, race, and ethnicity. Only images depicting a full face from a front perspective were selected, and the face part was cropped for training and testing. We applied standardized computer vision techniques using OpenCV (https://opencv.org/) for image reprocessing. Specifically, the computer algorithm recognized the human face in each image and cropped it with a bounding box to maximize useful information, which contained more percentages of facial features information. During the training, four images of each person were used for training, and one was reserved for testing. The full list of images can be found in the **Supplementary Materials**. After experimenting with various manipulations, we will use 200-by-200 pixels in this study as they provide the optimal balance between time efficiency and model effectiveness. Therefore, all images were scaled to a 200-by-200 pixel size including the face area from the original images.

### Training and testing procedures

The manipulation of E/I imbalance in CNN models. The E/I imbalance theory posits that there is an excessive excitatory signal in the neural systems in ASD^17^, and therefore, we simulated the excitability of the CNN models by manipulating the slope of the activation function, namely, the ReLU function (**Equation 1.1**).

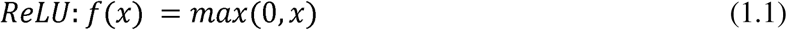

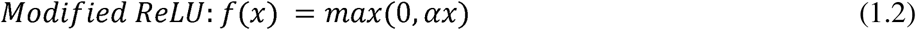

In our simulations, we manipulated the *α* value in the modified ReLU function (**Equation 1.2**) in the convolutional layer, a scaling factor that controls the output of the positive values of all units. Conventionally, the *α* value was set to 1. Through pilot testing, we found that the CNN models with *α* = 0.05 represented the optimal learning outcomes, so we chose it as a typical level of excitation (i.e., an optimal balance between excitatory and inhibitory signals). For the over-excitation condition, we set the *α* value to 0.5, ten times larger than the optimal *α* value. We also included an under-excitation condition as a comparison, in which the *α* value was set to 0.005, one-tenth of the optimal value (**Figure 1B**). Note that we did not aim to argue for a numeric equivalence of the over- and under-excitation conditions between the CNN model simulations and the biological conditions in humans. These values were chosen to merely demonstrate the impact of a single parameter that bears neurobiological implications to the E/I imbalance theory. We hypothesized that an over-excitation in CNN models would hinder the face recognition ability by weakening the pattern separation in the internal neural representations, whereas the under-excitation would also affect the learning of faces by forcing pattern separation excessively.

The manipulation of Internal Noise (IN) in CNN models. To test the predictions of the Internal Noise (IN) theory in CNN models, we introduced a Gaussian noise layer after each Convolutional Block (**Figure 1C**), adding intrinsic but random noises into the signals after the dropout layers and before the next processing layers. In this way, the output of each unit from the dropout layers was offset by a value randomly sampled from a Gaussian distribution with a standard deviation of *σ* (**Equation 3**) and a mean of 0. For the typical condition, we set the *σ* = 0 as in a system with minimal disturbance. Then, we changed *σ* to 0.5 (a mildly noised condition) or 5 (a severely noised condition) to examine the influence of the additional internal noise on learning accuracy as well as the pattern separation in the internal representations.

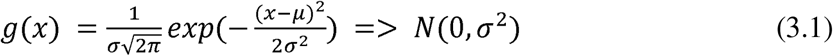

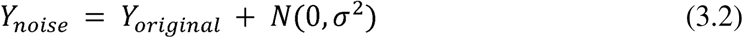

General training and testing settings. We ran all the CNN models for 1000 epochs, and for each condition (i.e., three conditions for the E/I and three for the IN simulations), we ran 20 individual loops with random initial weights to represent variabilities across individuals with the same training and testing procedures^40^. All models were trained with the Adam optimizer^50^ with a learning rate of 0.001 and binary cross-entropy loss function. For each training epoch, a batch size of 64 was used with standardized settings. Different images of the same individuals were trained to be associated with the same output label, whereas different output labels were used for different individuals in a localizationist manner. The output layer utilized the SoftMax function to generate the final label according to each input stimulus pattern (i.e., each image).

Scoring face recognition accuracy. Since we were mostly interested in how different parameters could lead to different learning performance and internal representations, we focused our analysis on trained items (i.e., four trained images of each individual). For each loop under each condition, we scored the accuracy by dividing the total number of corrected predictions by the total number of predictions (i.e., 4 X 10 = 40 total predictions; **Equation 4**) at each training epoch. Then, we compared the mean accuracy of the different loops between different conditions of E/I or IN simulations, to test their hypotheses on learning performance of face recognition. When examining the effect over training, the accuracy was sampled every 10 epochs.

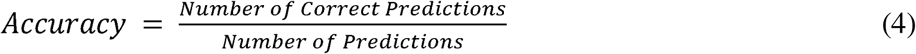

Representational Similarity Analysis. Following the standard practice in relevant computational and neuroimaging research^33,51–53^, we generated the similarity matrix by calculating the Pearson’s *r* between every pair of trained images (4 images for each of the 10 individuals) over the units in the flattened layer. We chose the flattened layer because it was the last layer before the output layer, so it contained the highest level of the internal representations of all trained images in the CNN models. We then computed the mean correlation for images of the same individual (within-person similarity) and the mean correlation for images of different individuals (between-person similarity). Success in face recognition requires an ability to recognize the same person across images, i.e., a high within-person similarity, and also the ability to differentiate images of different individuals, i.e., a low between-person similarity. These measures could provide insights on how the CNN models generalize across images for the same individuals, and different images of different individuals. Furthermore, comparing the within- and between-person similarities across different conditions of E/I or IN conditions, we could assess the impact of E/I and IN on the ability to represent face stimuli within the CNN models.

After the models were trained, we fed all images (including training and testing sets) and extracted the activation values of all units in the flattened layer. We then calculated the Pearson correlation between all possible pairs of the images based on the activation values of 64 units in the flattened layer. The similarity matrices of each loop after 500, 750, and 1,000 epochs were computed, and the mean within- and between-person similarity was calculated separately. A high similarity value suggests overlapping representations whereas a low similarity value suggests separable representations in the CNN models.

## Result

### Impaired learning ability due to E/I imbalance and internal noise

**Figure 2A** shows the training accuracy of different conditions of E/I imbalance (i.e., the slope of the ReLU function). A significant main effect of E/I imbalance was observed over the training, *F*(2, 5757) = 23051.581, *p* < 0.001, *η*^2^ = 0.889. There was also a significant interaction between Slope and Epoch, *F*(200, 5757) = 18.258, *p* < 0.001, *η*^2^ = 0.388. Overall, the pattern stabilized after 300 epochs of training. The over-excitation condition (ReLU slope = 0.5) failed to learn the trained images of faces compared to the other two conditions, even after 500 epochs, *F*(2,57) = 225.25, *p* <0.001, *η*^2^ = 0.888, while the other two conditions almost excelled at their performances (see **Table 1**). The under-excitation condition (ReLU slope = 0.005) showed a slight disadvantage in learning at the early stage (e.g., 100 epochs) compared to the optimal E/I condition, *t*(38) = −5.47, *p* = 4.98×10^−5^ (Tukey adjusted), and quickly caught up in the performance and eventually achieved a comparable accuracy at 300 epochs (HSD adjusted *p* = 0.905).

**Figure 2.**
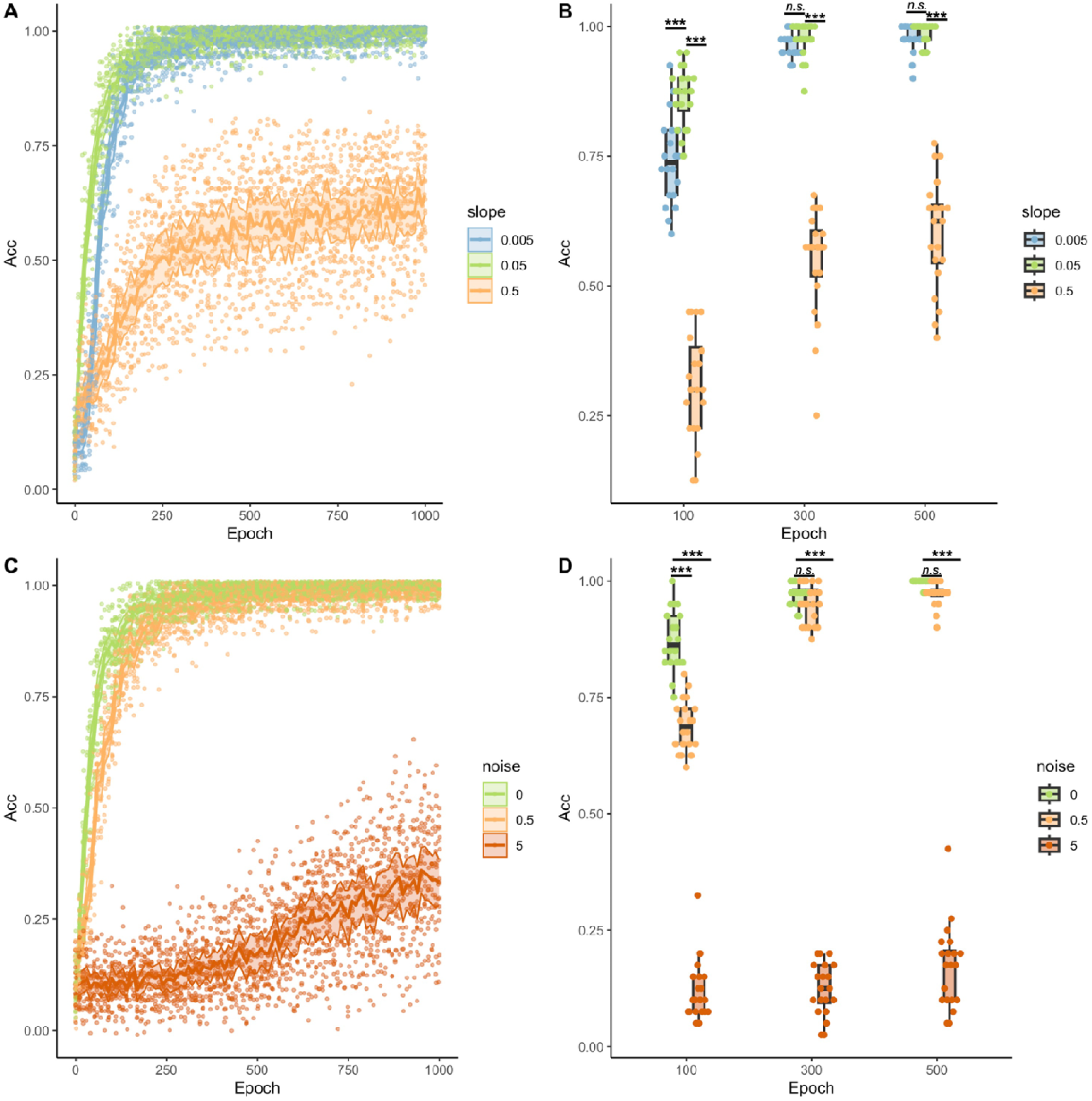
Accuracy of face recognition in CNN models on trained dataset. (**A**) The face recognition accuracy of models with different slopes in ReLU function of the convolutional layers (0.05 = optimal; 0.005 = under-excitation; 0.5 = over-excitation); (**B**) The face recognition accuracy of CNN models with different strengths of noises added to the weights (the standardized deviation of the Gaussian noise: 0 = optimal; 0.5 = mild; 0.5 = over-excitation). In A and B, accuracy was assessed every 10 epochs (shown in individual dots), and the solid lines showed the group average over 20 different runs. The Shades around the solid lines depicted the 95% confidence interval. (**C**) and (**D**) showed the accuracy at selected epochs to demonstrate the performance differences over training in boxplots. *** <0.001.

**Figure 3.**
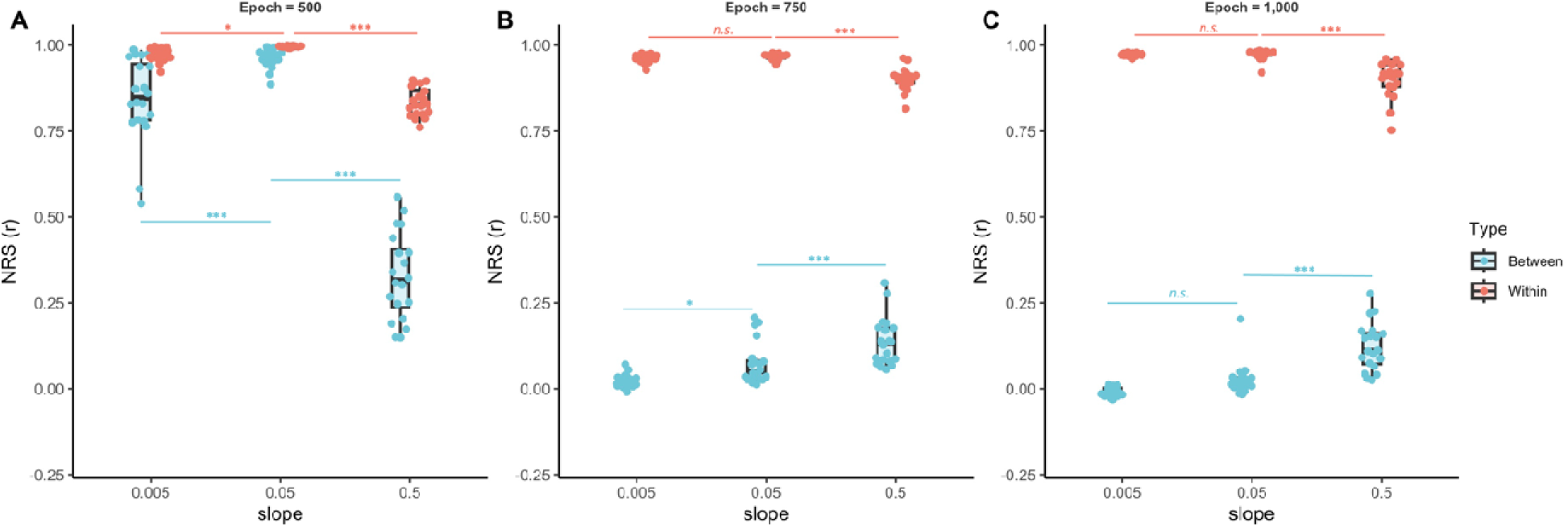
Neural representational similarity (NRS) in the last dense layer of CNN models with different manipulations of the slope in the ReLU function (for E/I imbalance) at (**A**) 500 epochs, (**B**) 750 epochs), and (**C**) 1,000 epochs. Blue boxes present the NRS of images of different person identities (between-identity NRS), and red boxes present the NRS of images of the same person (within-identity NRS). The optimal learning should show high within-identity NRS (i.e., overlapping units to represent different images of the same person in the model) and low between-identity NRS (i.e., separate sets of units to represent images of different persons). *<0.05, **<0.01, and ***<0.001.

**Table 1.** Descriptive data and ANOVA results of performance accuracy in face recognition.

| *ReLU Slope* |  |  |  |  |  |  |
| --- | --- | --- | --- | --- | --- | --- |
| Epoch | Optimal (0.05) | Under-excitation (0.005) | Over-excitation (0.5) | ANOVA | HSD test (0.005 vs 0.05) | HSD test (0.5 vs 0.05) |
| 100 | 0.87 (0.06) | 0.74 (0.08) | 0.31 (0.10) | $F(2,57) = 243.071$,<br>$p = 1.25 \times 10^{-28}$, $\eta^2 = 0.895$ | -0.125*** | -0.559*** |
| 300 | 0.98 (0.03) | 0.97 (0.02) | 0.55 (0.11) | $F(2,57) = 281.838$,<br>$p = 2.79 \times 10^{-30}$, $\eta^2 = 0.908$ | -0.009 | -0.477*** |
| 500 | 0.99 (0.02) | 0.98 (0.03) | 0.6 (0.11) | $F(2,57) = 225.25$,<br>$p = 8.66 \times 10^{-28}$, $\eta^2 = 0.888$ | -0.008 | -0.385*** |
| *Gaussian Noise* |  |  |  |  |  |  |
| Epoch | Optimal (0) | Mild (0.5) | High (5) | ANOVA | HSD test (0.5 vs 0) | HSD test (5 vs 0) |
| 100 | 0.87 (0.06) | 0.69 (0.06) | 0.12 (0.06) | $F(2,57) = 824.479$,<br>$p = 8.54 \times 10^{-43}$, $\eta^2 = 0.967$ | -0.184*** | -0.755** |
| 300 | 0.97 (0.02) | 0.95 (0.04) | 0.12 (0.06) | $F(2,57) = 2560.188$,<br>$p = 1.55 \times 10^{-56}$, $\eta^2 = 0.989$ | -0.02 | -0.845*** |
| 500 | 1.00 (0.01) | 0.97 (0.03) | 0.17 (0.09) | $F(2,57) = 1511.989$,<br>$p = 4.12 \times 10^{-50}$, $\eta^2 = 0.981$ | -0.025 | -0.825*** |

Notes: HSD post-hoc tests were done to compare the other conditions to the optimal condition, and the adjusted $p$ values are reported here. \*\*\*<0.001.

A similar pattern was observed for the manipulation of Internal Noise. The main effect of the Gaussian noise (no noise, mild noise with an SD = 0.5, high noise with an SD = 5) was significant, *F*(2,5757) = 100869.373, *p* <0.001, *η*^2^ = 0.972, suggesting that an increased level of random noises on the weights between layers led to compromised learning performance. The interaction between Noise and Epoch was also significant, *F*(200,5757) = 45.897, *p* <0.001, *η*^2^ = 0.615. Over the training, models with high noise consistently underperformed compared to the no noise and mild noise conditions (see **Table 1**). However, the difference between the no noise and mild noise condition was only significant at the early stage (100 epochs of training), *t*(38) = - 9.687, *p* = 1.20×10^−11^ (HSD adjusted), but became insignificant after 300 epochs of training (Tukey adjusted *p* = 0.315).

Our simulations showed that over-excitation or high level of internal noise in a learning system led to compromised ability for face recognition, as predicted by the E/I imbalance theory and internal noise theory. However, the impact of under-excitation or mild level of internal noise only had an impact on learning during the early stage, but not necessarily disrupted the learning in the long run.

### Atypical neural representation due to E/I imbalance and internal noise

Next, we examined the neural representation similarity (NRS) between images, and we specially investigated the NRS of images of the same person identity (within-person) vs. the NRS of images from different person identities (between-person). Although the learning accuracy was stable after 500 epochs of training for both simulations, it was surprising to observe that within- vs. between-person NRS were not quite separate from each other for models with optimal settings (i.e., slope = 0.05 and Gaussian noise = 0). Therefore, we examined the change of NRS from 500, 750, and 1,000 training epochs.

The NRS matrices of all trained images were shown in **Figure S1** for the manipulation of the ReLU slope. Overall, the within-person NRS values were higher than between-person NRS values across different training epochs, but the separation became larger with more training. When the model was over-excited (i.e., slope = 0.5), the between-person NRS seemed to be high even after the model was trained for 1,000 epochs. A mixed ANOVA with Slope (0.005, 0.05, and 0.5) as a between-subject factor and with NRS Type (Within vs. Between) as a within-subject factor revealed significant main effects of both Slope and Type for all three training epochs (see **Table 2**). At 500 epochs, the optimal slope (i.e., 0.05) showed the highest within-person NRS compared to the other two conditions (both *p*s < 0.05, HSD adjusted), but also the highest between-person NRS (both ps < 0.01, HSD adjusted). In contrast, after 750 epochs of training as well as seen for 1,000 epochs of training, the over-excitation models demonstrated the lowest within-person NRS but the highest between-person NRS compared to the optimal and under-excitation conditions, suggesting a weak ability to represent images of the same person in a similar way but also to differentiate images of different persons at the representational level.

**Table 2.**
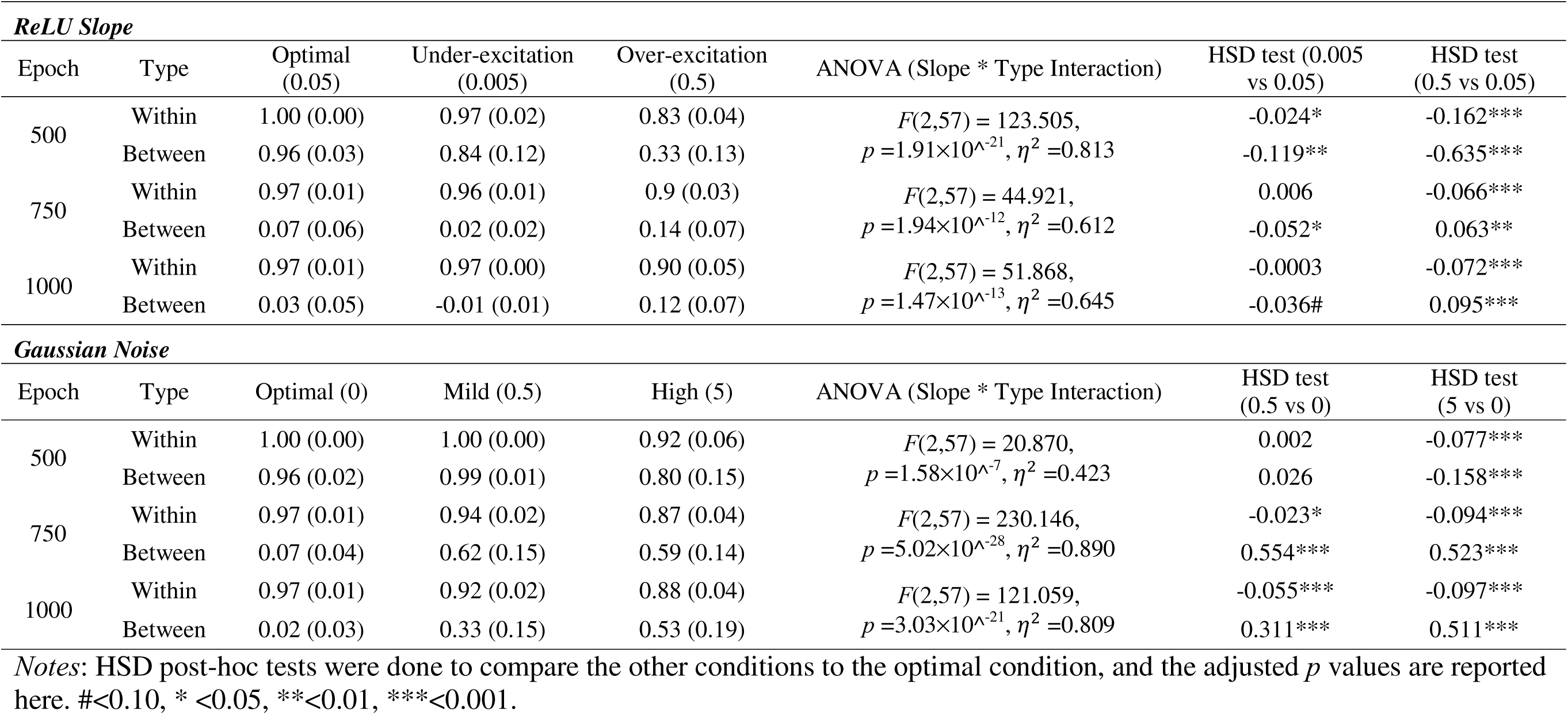
Descriptive and ANOVA results of within-person and Between-person neural representation similarity (NRS).

The results for NRS with noise manipulation showed a similar pattern (NRS matrices in **Figure S2**): both within- and between-person NRS values were high after 500 epochs of training, and the separation was evident with more training. At 750 epochs, the optimal condition (noise SD = 0) showed the highest within-person NRS with the lowest between-person NRS, whereas the results for models with moderate or severe noise (SD = 0.5 or 5) still demonstrated relatively high levels of between-person NRS (see **Table 2**). At 1,000 epochs, there was a graded effect that the larger the noises applied to the weights, the less separation we observed for the within- and between-person NRS values. This finding suggests that increased internal noises in a learning system also led to weak representations of face images.

### Diffused pattern representation due to E/I imbalance and internal noise

To further explore how the E/I imbalance and internal noise impact the neural representations, we calculated the percentage of inactive units in the convolutional layer of all three convolutional blocks (before normalization, pooling, and dropout layers), as an index of sparseness. A unit was considered inactive if its activation value was below 0.001, often appearing black in visualized feature maps. The assumption is that a few units are engaged or activated when the model efficiently represents the stimuli. For E/I imbalance, simulations with the optimal slope showed a high rate of inactive units across three layers, especially compared to the over-excitation condition (all *p*s < 0.001; see **Table 3**). The under-excitation condition showed low levels of sparse representations at the first two layers but a high rate of inactive units at the 3rd convolutional layer, even significantly higher than the optimal slope condition (**Figure 4**; and also in **Figure S3**). For internal noise, the optimal condition (no noise) unsurprisingly showed the highest rate of inactive units across all layers (**Table 3 and Figure 4**), with the mild noise condition showing the lowest rate overall. Therefore, our results suggest that the optimal conditions for both E/I imbalance and internal noise stimulations not only represent individual faces effectively (i.e., high within-person but low between-person NRS), but also efficiently (i.e., with fewer units being activated).

**Table 3.** Descriptive and ANOVA results of inactive units in convolutional layers.

| ReLU Slope |  |  |  |  |  |  |
| --- | --- | --- | --- | --- | --- | --- |
| Layer | Optimal (0.05) | Under-excitation (0.005) | Over-excitation (0.5) | ANOVA (Slope * Layer Interaction) | HSD test (0.005 vs 0.05) | HSD test (0.5 vs 0.05) |
| Conv1 | 75.34 (5.71) | 48.38 (9.57) | 49.30 (12.92) | $F(4,171) = 34.508,$<br>$p = 4.16 \times 10^{-21}, \eta^2 = 0.447$ | -27.00*** | -26.00*** |
| Conv2 | 73.66 (4.87) | 53.03 (2.74) | 50.02 (0.84) |  | -20.60*** | -23.60*** |
| Conv3 | 70.56 (7.09) | 75.14 (4.15) | 52.68 (1.68) |  | 4.58* | -17.90*** |
| Gaussian Noise |  |  |  |  |  |  |
| Layer | Optimal (0) | Mild (0.5) | High (5) | ANOVA (Slope * Layer Interaction) | HSD test (0.5 vs 0) | HSD test (5 vs 0) |
| Conv1 | 76.08 (5.1) | 47.6 (8.7) | 63.18 (11.09) | $F(4,171) = 3.614,$<br>$p = 0.007, \eta^2 = 0.078$ | -28.50*** | -12.90*** |
| Conv2 | 74.06 (5.01) | 48.75 (1.21) | 55.73 (1.64) |  | -25.30*** | -18.30*** |
| Conv3 | 70.8 (6.15) | 49.17 (7.96) | 55.09 (6.08) |  | -21.60*** | -15.70*** |

Notes: HSD post-hoc tests were done to compare the other conditions to the optimal condition, and the adjusted $p$ values are reported here. \* <0.05, \*\*\*<0.001.

**Figure 4.**
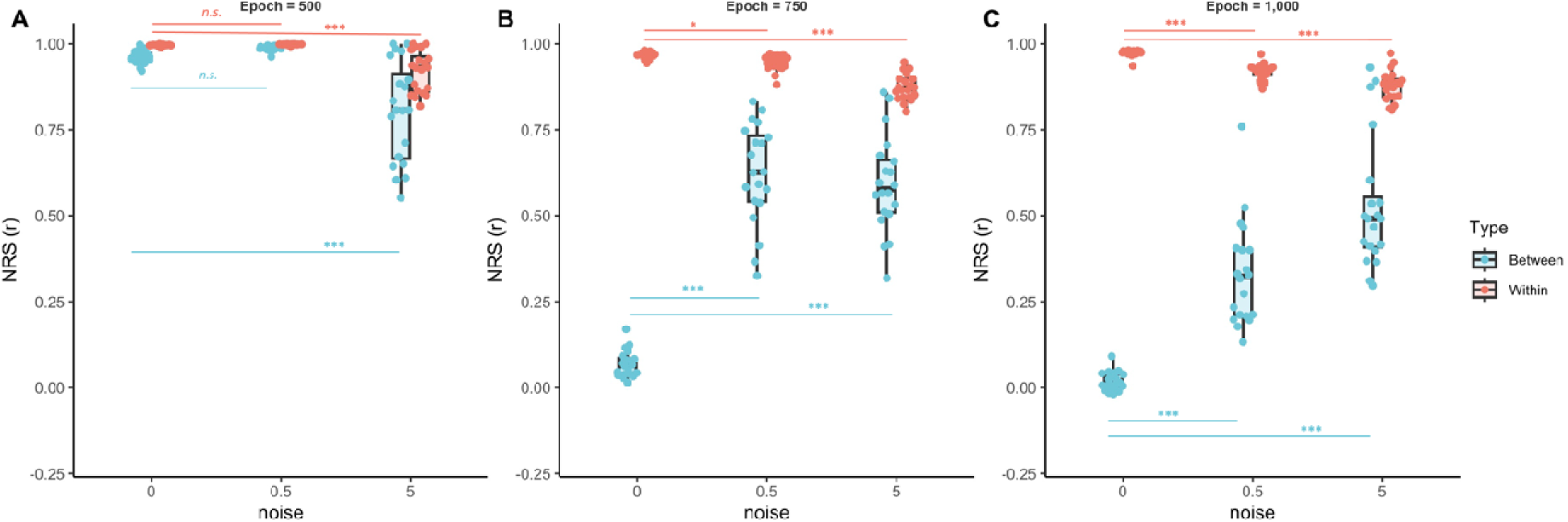
Neural representational similarity (NRS) in the last dense layer of CNN models with different manipulations of the Gaussian noises added to the weights (for internal noise) at (**A**) 500 epochs, (**B**) 750 epochs), and (**C**) 1,000 epochs.

**Figure 5.**
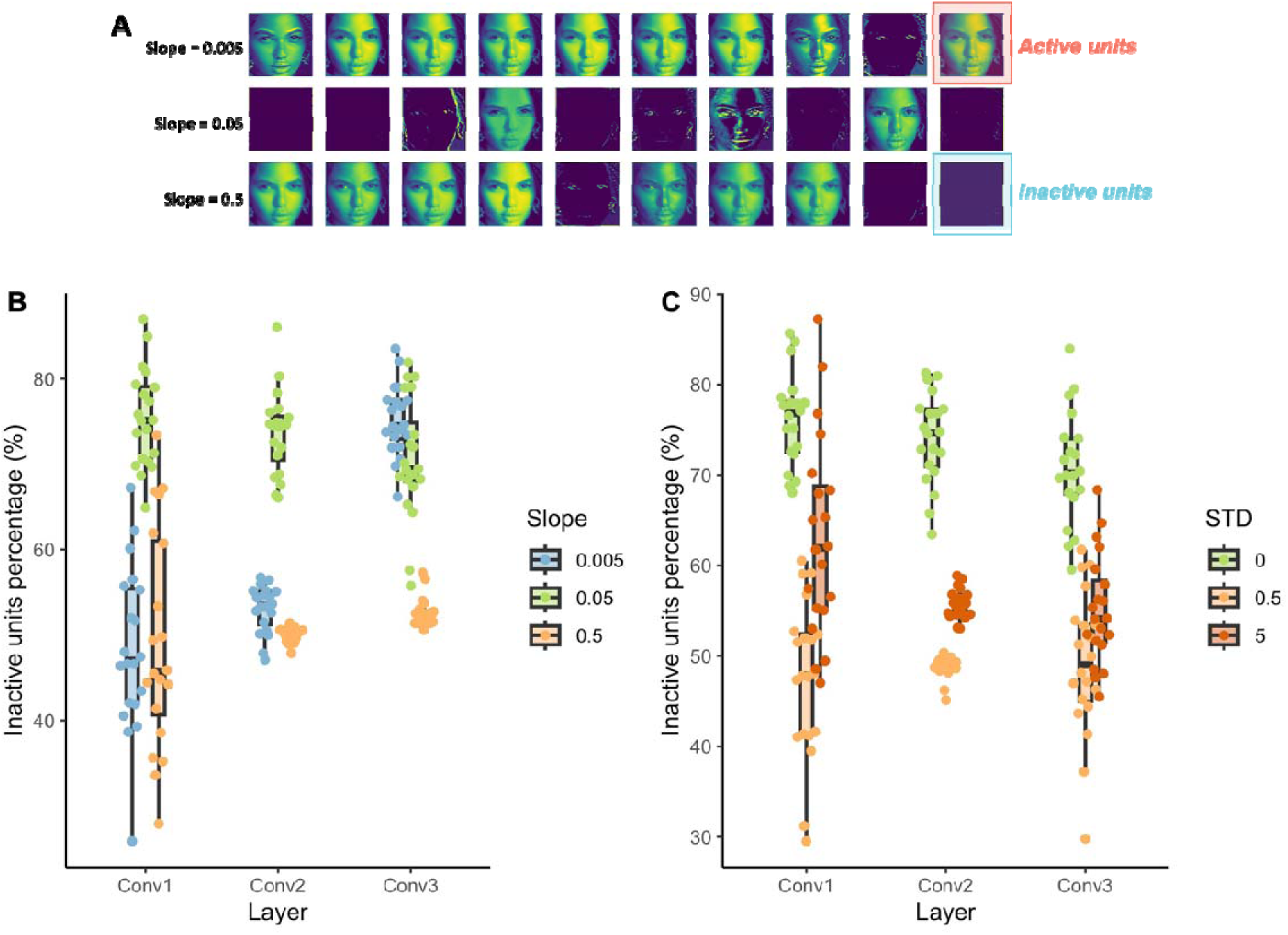
The percentage of inactive units in the convolutional layers after 1,000 training epochs. (**A**) A sample demonstration of face representation in the first convolutional layer of models with different slopes in the ReLU function. (**B**) and (**C**) show the percentage of inactive units across three convolutional layers for simulations of the effect of ReLU slope (i.e., E/I imbalance; B) and Gaussian noise (i.e., internal noise; C).

## Discussion

Our findings suggested that both E/I imbalance and internal noise were associated with reduced ability in face recognition in CNN models. Specifically, under-excitation led to inferior face recognition early during the learning, but over-excitation led to a sustaining deficit in face recognition. However, increased internal noises were associated with impaired face recognition in a graded way: the larger the noise was, the worse the learning was. For internal representations, it was shown that over-excitation resulted in slightly reduced within-individual similarity and increased between-individual similarity, likely resulting from excessive activations in a high percentage of CNN units. By contrast, the increased internal noise led to a monotonic change with decreased within-individual similarity but increased between-individual similarity, associated with a higher percentage of excessively activated CNN units.

The results based on E/I imbalance simulations were largely consistent with the predictions and existing literature. As predicted by the E/I imbalance theory, ASD is associated with excessive excitatory signals in the neural system, leading to cognitive challenges^17,19,20^. In our E/I imbalance simulations, we changed the slope of the ReLU function to approximate the excessive excitatory signals in the human brain^36^. We showed that the over-excitation led to delayed learning of face identity and reached the plateau after 500 training epochs. This is consistent with the growing body of literature that autistic individuals have difficulties in face recognition^2,3,7,23,51,52^ and even episodic memory in general^53–57^. Our simulations further suggested that over-excitation in CNN models led to atypical neural representations of faces compared to the optimal condition, particularly for the faces of different individuals, i.e., elevated levels of between-person similarity, as well as a slight decrease in within-person similarity, i.e., faces of the same individual. Only a handful of studies examined neural representation similarity in ASD so far^33,61–64^, and a couple of studies showed that autistic individuals failed to show differential neural representations for different types of social intents^64^ or functions^62^. One study examined the neural representational similarity in the right fusiform face area (FFA) for face processing, and showed lower neural representation similarity of the same kind of objects (faces or cars) in autistic adults compared to non-ASD^33^. Therefore, our results replicated these findings, and elucidated that reduced face recognition ability can be due to over-excitation and an inability to establish differential neural representations of faces from different individuals. These results are probably due to unnecessarily activated units in the different CNN layers, leading to using many overlapping units to represent the faces of different individuals (i.e., lack of sparse representations). This finding is also consistent with the recent study suggesting that a subgroup of autistic children showed atypical pattern separation memory for daily objects with more diffused memory representations for similar memory experiences^66^.

Interestingly, the under-excitation condition showed mild influence on face recognition only at the early stage of learning and even more differentiated representations of the faces of different individuals (at 500 epochs, optimal: 0.96 vs under-excitation: 0.84; p < 0.01). Thus, over- and under-excitation seemingly led to opposite patterns of neural representations, which is consistent with the recent findings that the pattern separation memory in ASD is heterogeneous, and there are subgroups with both over-differentiated and under-differentiated representations of memory experience^66^. This pattern potentially represents a large subgroup of autistic individuals who have more isolated memory experiences and a lack of generalization ability across similar experiences^66,67^. Thus, the predictions from E\I imbalance were largely supported by our simulations and consistent with the literature on face recognition in ASD. More importantly, our simulation based on E/I imbalance could potentially explain the different profiles of atypical face recognition and memory representations in ASD.

The results from IN simulations were also consistent with the hypothesis that increased noises will lead to lower face recognition ability^18,29,70^. However, instead of observing reduced representation similarities overall, we found that the increased noise weakened the representations of faces of the same individual, i.e., less similar, but also confused the representations of faces of different individuals (i.e., more diffused representations). This finding is largely similar to the E/I simulations with over-excitations and contradictory to our hypothesis. One possible explanation is that the random noises added into the learning system led to random activations in CNN units at the early stage of learning. Once these units were activated, they were utilized to process the faces of different individuals, leading to overlapping representations of the faces of different individuals on these randomly activated units. At the same time, the randomly activated units also led to less overlapping representations of the faces of different individuals since they were unnecessary to represent the faces of the same individual. For this reason, we observed a high percentage of activated CNN units with both mildly- and severely-noised conditions for face presentations. Our IN and E/I (over-excitation) simulations seem to suggest the intrinsic link between the theories of E/I imbalance^17^ and internal noise^18^. The large internal noise in the neural system of ASD may simply be an epiphenomenon of excessive excitatory signals. If so, although these two theories may capture different features of the neurobiological dysfunctions of ASD, they may be two sides of the same coin. However, based on our simulation, the effect of the internal noise was rather monotonic that weaker/more diffused representations were found with increased noises. Unlike the results from E/I imbalance, it failed to explain the full spectrum of atypical memory abilities and representations observed with both over- and under-differentiated memory representations in ASD revealed in recent studies^65,67^.

Using a computational approach, our studies suggested that E/I imbalance and increased internal noises in the neural system could underlie the face recognition challenges in autistic individuals. Using a theory-driven approach, we were able to demonstrate the utility and power of using computational models to understand human cognition and neurocognitive mechanisms of cognitive disorders^38,71,72^, as well as its ability to bridge the behavioral and neuroimaging studies to examine the individual differences^37,40,73,74^. However, we would emphasize again that by no means do we argue for a neurobiological faithfulness of these simulations and parameters in CNN models. In fact, we only consider this theory-driven approach a tool to test predictions of theories in a well-controlled learning system that can hardly be done in human research. We need converging evidence from behavioral, neuroimaging, and computational approaches to establish, test, and revise key theories of human cognition and to understand the neurobiological mechanisms of cognitive challenges in clinical populations such as ASD. We believe that this integrated approach can be easily applied to other domains to advance our knowledge of other types of developmental and neurobiological disorders as well as their cognitive challenges. Moreover, we also do not argue that these two neurobiological factors are the only reasons why face recognition is challenging in ASD. These theories may only explain certain subtypes of face recognition challenges in ASD. Other possibilities, such as lack of social motivation^4,75^ or dysfunctions in the rewarding system for social stimuli^76,77^, could still result in face recognition challenges in ASD with seemingly similar behavioral outcomes, but different underlying mechanisms such as hypo-reactivity or hypo-connectivity within the extended face processing network^31,76^. Future work should examine how these factors interact and jointly affect the different phenotypes of ASD and their heterogeneous presentations of cognitive and social challenges^77–81^.

In sum, our computational simulations demonstrated that both the E/I imbalance and internal noise (IN) theories can explain the variability in face recognition ability in ASD. In addition, the CNN models allowed us to link the impaired behaviors to internal representations as well as neurobiological theories^37,47,52^. Such an approach provides an ideal vehicle to integrate neurobiological, behavioral, neuroimaging, and computational research to further our understanding of neurological and psychiatric disorders^78,82^. This generalizable framework can be easily applied to other clinical disorders and potentially sheds light on clinical and educational practices.

## Acknowledgments

We thank the Santa Clara University for providing WAVE Visualization internal grant to L.C. as well as a REAL summer scholarship to X.W. We also thank Akshat Karla for his early contribution to this project.

## Data and code availability statement

All the data and analytical codes can be downloaded from https://github.com/xthomaswang/ASD_FaceReg_Modeling_CNN.

## Competing interests

All authors claim no conflict of interest.

## Notes

https://github.com/xthomaswang/ASD_FaceReg_Modeling_CNN

### Competing Interest Statement

The authors have declared no competing interest.

