## Supplementary Information for "A neurocomputational basis of face recognition changes in ASD: E/I balance, internal noise, and weak neural representations"

Wang et al.

*Table of contents*

1.1 Supplementary materials
1.2 Supplementary figures

**Supplemental Materials**

All cropped images used for the model simulations are listed below. One of the five images of the same person was reserved as a testing image for the identity.

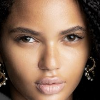

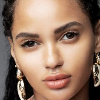

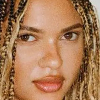

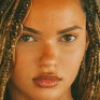

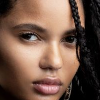

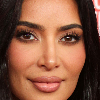

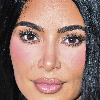

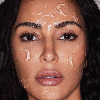

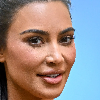

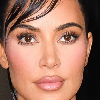

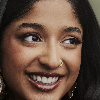

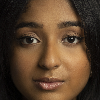

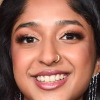

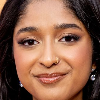

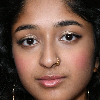

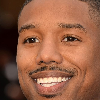

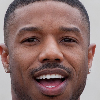

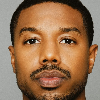

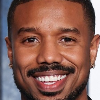

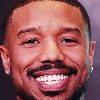

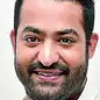

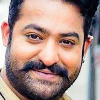

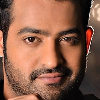

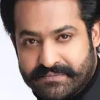

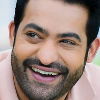

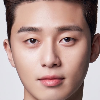

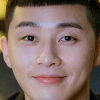

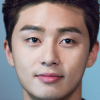

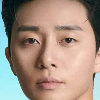

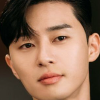

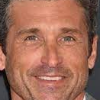

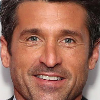

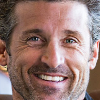

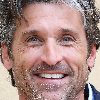

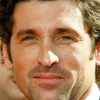

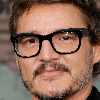

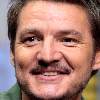

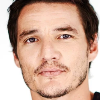

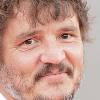

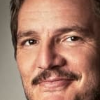

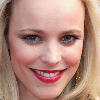

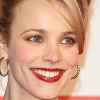

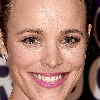

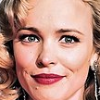

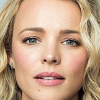

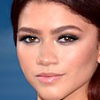

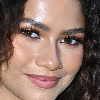

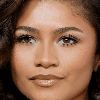

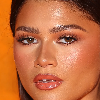

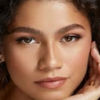

**Supplemental Figures**

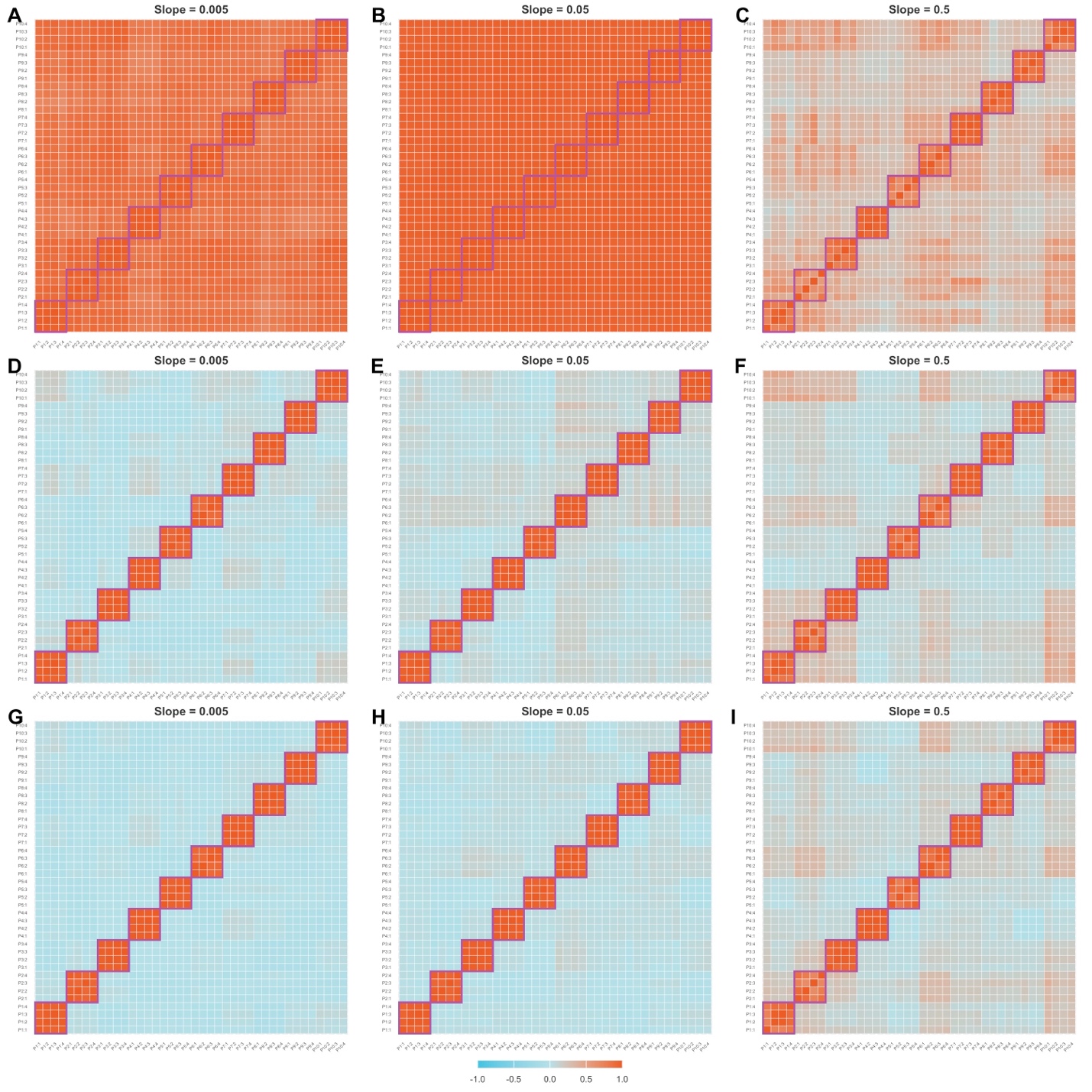

**Figure S1**. Similarity matrices for all trained images with the E/I imbalance simulations (i.e., manipulations of the ReLU slope). (**A-C**) for 500 training epochs, (**D-F**) for 750 training epochs, and (**G-I**) for 1,000 training epochs. The rows and columns of the matrices were organized by the trained images and for example, P1:1 stands for image 1 for person 1. Therefore, the diagonal squares with high correlation values (close to 1) suggest high similarity between different images of the same person (bordered by the purple lines).

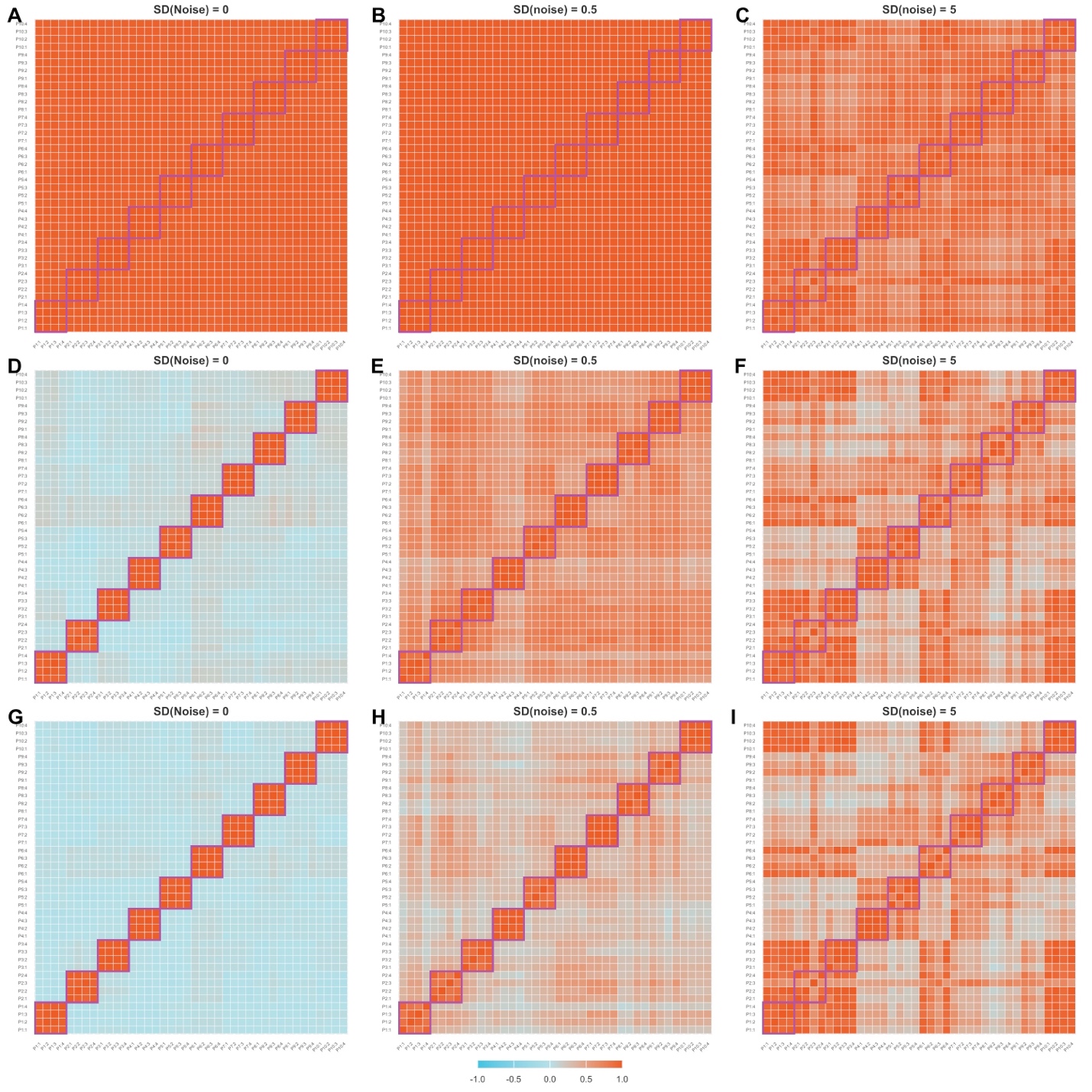

**Figure S2**. Similarity matrices for all trained images with the internal noise imbalance simulations. The setups are similar to those in **Figure S1**.

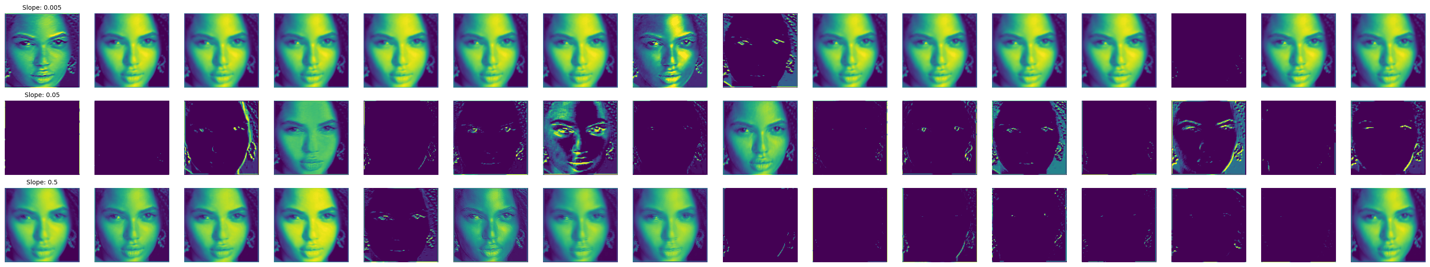

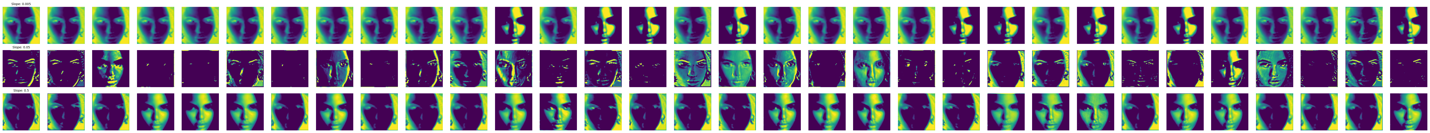

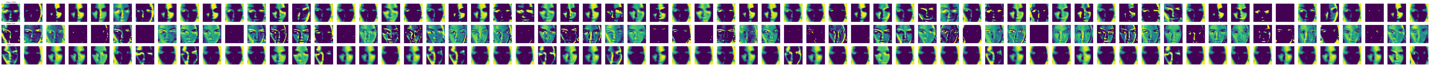

**Figure S3**. Samples of unit activation patterns of the convolutional layers when the slope was set to 0.005 for Image 1 of Person 1. From the top to the bottom, it shows convolutional layers 1-3.
